# Cascading periods of language-related brain plasticity across early childhood

**DOI:** 10.64898/2026.03.27.714739

**Authors:** Monica E. Ellwood-Lowe, Monami Nishio, Alexander J. Dufford, Michael Arcaro, Theodore D. Satterthwaite, Allyson P. Mackey

## Abstract

Language is thought to have multiple sensitive periods in early childhood, but the neural basis of these sensitive periods is less understood. We leverage advances in in-vivo neuroimaging of plasticity, measuring the neural inhibition across the brain via Hurst exponent. Using two large datasets with children ages 10 months to 18 years (Baby Connectome Project: 10m-3y6m, 458 observations across *n*=222 children; Human Connectome Project-Development: 5-18y, *n*=437), we characterize the development of the Hurst exponent in language-related brain regions. In early childhood, Hurst increases in temporal and frontal language areas, and posterior regions develop earlier than anterior regions. In contrast, thalamic Hurst plateaus earlier, perhaps underlying the earliest language-related sensitive periods. Children with higher language-related skills show slower increases in cortical Hurst in early childhood, suggesting protracted plasticity. Later in childhood, cortical Hurst plateaus around age 9, suggesting a potential neural mechanism for age-related declines syntax learning. These results highlight a potential neural basis for cascading language-related sensitive periods.

**Research highlights:**

- Language has multiple, cascading sensitive periods, but the neural basis of these sensitive periods is not well-understood.
- We leverage advances in in-vivo neuroimaging to quantify plasticity in language-related brain regions across childhood via Hurst, a measure of inhibition.
- We find that Hurst increases (plasticity decreases) in a graded fashion, with posterior regions maturing before anterior regions.
- Thalamic hurst plateaus in the first year of life, while cortical Hurst plateaus at age 9, suggesting a neural basis for distinct language-related sensitive periods.

## Introduction

Learning a new language is difficult for adults, but remarkably easy for children. This has long been explained by sensitive periods in language development: the idea that there are moments in development when children’s brains are particularly receptive to linguistic information, after which plasticity diminishes.^1^ Behaviorally, there is now ample evidence that there is not just one language-sensitive period, but a cascading set of sensitive periods for different language-related skills (e.g., phonetic perception, word pronunciation, syntax).^1^ However, our understanding of the neural mechanisms underlying these sensitive periods is still sparse. Understanding the neural basis for sensitive periods in language development has the potential to provide foundational insights into basic questions about the neural basis of language as well as applied questions about when and how it becomes harder to learn a new language.

For hearing children, learning to distinguish speech sounds (phoneme perception) scaffolds later semantic and syntactic learning, each of which may have its own sensitive period. Phoneme perception is perhaps the best-characterized example of a sensitive period in language development, with children losing the ability to hear speech sounds not used in their native language between 6-12 months of age.^2–4^ For example, Hindi distinguishes between *ṭa* and *ta*, a contrast that non-Hindi speakers lose the ability to hear. Similarly, English distinguishes between *ra* and *la*, which Japanese does not use, and non-English-speaking Japanese speakers lose the ability to hear.^2,5^ Experience hearing a language in infancy seems to be sufficient to retain sensitivity to that language’s phonetic distinctions, even in the absence of later exposure.^6,7^

Later in infancy, children show a rapid increase in word comprehension, followed by a stereotyped spurt in word production between ages 18-24 months. Next, they develop language-specific syntactic rules, which are another aspect of language thought to be governed by sensitive periods. While there is mixed evidence for the timing of sensitive periods in syntax, evidence from international adoptees and second-language learning has generally pointed to a sensitive period around age 8,^8,9^ though other evidence points to a sensitive period much later, around age 15-17.^10,11^ Thus, the evidence to date suggests that peak plasticity for syntax appears to be reached by age 8 at the earliest, or age 17 at the latest. Taken together, the language system follows from a set of cascading sensitive periods, beginning in the first year of life and spanning later childhood to adolescence.

At the neural level, language relies on a distributed set of regions across left superior temporal and frontal cortex, spanning lower-order sensorimotor cortex to higher-order association cortex.^12,13^ The neural system for language appears to be in place by early childhood, though the language system becomes increasingly sensitive to language across this period.^14^ The specific role of distinct language-related brain regions is contested. Classic accounts suggest temporal regions are more involved in comprehension, while frontal regions are more involved in production.^15^ In support of this possibility, early studies found that people with damage to left superior temporal areas (Wernicke’s area) are more likely to show deficits in comprehension, while those with left inferior frontal areas show deficits in production.^15^ However, large-scale evidence suggests that at least in adulthood, all of these regions work in concert to perform various language tasks.^16–19^ Whether the same is true in childhood, as children are acquiring language, remains less clear.

The thalamic nuclei also play a central role in the language system. Left thalamic nuclei are consistently activated along with frontal and temporal language areas in fMRI studies.^13,20–22^ Supporting its causal role, deep-brain stimulation of thalamic nuclei in adult neurosurgery patients affects both semantic and syntactic performance.^23^ The medial geniculate nucleus (MGN) is connected to the primary auditory cortex and is thought to play a role in the early gating of sound information, which may be relevant for speech sound processing.^24,25^ Indeed, there is evidence that MGN plays an important role in processing phoneme distinctions.^26–28^ On the other hand, the ventral lateral nuclei (VL) play a role in language production and show white matter projections to the inferior frontal and motor cortex.^29,30^

Despite decades of work characterizing both milestones in language development and the neural basis of language, the neural mechanism underlying language-related sensitive periods remains unclear. One possibility is that the hierarchical development of the brain may underlie the cascading sensitive periods seen behaviorally. For example, sensorimotor regions of the brain develop earlier than higher-order association areas.^31^ A developmental precursor to the sensorimotor-association (S-A) axis may be the anterior-posterior axis^32^, in which posterior regions (toward the back of the brain) develop earlier than anterior regions (toward the front).^33,34^ Earlier-developing posterior temporal regions may align with the timing of earlier-developing language milestones (e.g., phoneme perception), while later-developing frontal regions may align with the timing of later language processes (e.g., production, syntax).

One of the reasons we have not clearly identified the neural basis of language-related sensitive periods is that brain plasticity in young children has been remarkably hard to measure *in vivo*. Recently, advances in technology have allowed us to infer neural inhibition—which underlies plasticity—via fMRI with young children.^35–37^ It is well established that increases in neural inhibition—particularly through the maturation of parvalbumin (PV) neurons—lead to reduced plasticity and the closure of sensitive periods.^38^ Therefore, tracking the development of inhibition allows us to quantify both the peak and the subsequent reduction of plasticity that define these critical developmental windows. One strong contender for measuring inhibition in vivo in humans is the Hurst exponent, which quantifies long-range temporal correlations and scale-invariant dynamics in various time series.^39^ Computational models have demonstrated that enhancing inhibition leads to an increase in the Hurst exponent of simulated local field potential and BOLD signals.^35,40–42^ Studies in rodents and non-human primates have shown that the Hurst exponent is sensitive to GABA changes, supporting a causal link between GABA and the Hurst exponent.^35,42^ Our recent work further supports the validity of the Hurst exponent as a developmental measure of inhibition by demonstrating spatial and temporal correspondences between the Hurst exponent and PV mRNA expression, as well as PV cell density, across species.^37^ Thus Hurst is a non-invasive way to identify the peak and subsequent plateau in inhibition that underlie sensitive periods *in vivo*.

Here, we used the Hurst exponent of fMRI data to examine the ages at which plasticity diminishes across language-related brain regions. We predicted that superior temporal regions would show a plateau in inhibition in the first year of life, in line with the timing of infants’ declining sensitivity to non-native phonetic categories. We posited that this plateau in inhibition might occur later in inferior frontal regions. We also predicted there would be cascading periods of inhibition across the sensorimotor-association gradient in language-related brain regions. Finally, we were interested in whether the trajectories of inhibitory development would be different depending on children’s early language skills. There is mixed evidence on whether brain development would be delayed as a result of more enriched input or alternatively sped up^31,43–45^, therefore, we considered both possibilities.

## Methods and Materials

### Participants

Data were drawn from two datasets: the Baby Connectome Project (BCP) and Human Connectome Project: Development (HCP-D). Detailed demographic information is provided in Table 1.

**Table 1.** Demographic Characteristics.

|  | **BCP (*N* = 222)**<br>**mean or #** | **HCPD (*N* = 437)**<br>**mean or #** |
| --- | --- | --- |
| **Age (years)** | 1.931 ± 0.6569 | 12.941 ± 2.810 |
| **Sex (male)** | 103 (46%) | 197 (45%) |
| **Ethnicity** |  |  |
| Hispanic or Latino | 14 (6%) | 63 (14%) |
| No Response | 0 (0%) | 4 (1%) |
| **Race** |  |  |
| White | 140 (63%) | 304 (70%) |
| Black | 4 (2%) | 29 (7%) |
| Asian | 1 (0%) | 18 (4%) |
| Others | 30 (14%) | 82 (19%) |
| No Response | 47 (21%) | 3 (1%) |
| **Income** |  |  |
| 25-35k | 7 (3%) | 9 (2%) |
| 35-50k | 3 (14%) | 22 (5%) |
| 50-75k | 36 (16%) | 40 (9%) |
| 75-100k | 32 (14%) | 57 (13%) |
| 100-150k | 56 (25%) | 107 (24%) |
| 150-200k | 27 (12%) | 72 (16%) |
| 200k- | 13 (6%) | 80 (18%) |
| No Response | 48 (22%) | 50 (11%) |
| **Parent Education** |  |  |
| Less than college | 13 (6%) | --- |
| College degree | 119 (54%) |  |
| Graduate degree | 42 (19%) |  |
| No Response | 48 (22%) |  |
| **Multilingual** |  | --- |
| Multilingual | 17 (13%) |  |
| Monolingual | 112 (50%) |  |
| No Response | 93 (37%) |  |

#### 1. BCP

Data were obtained from the Baby Connectome Project (https://nda.nih.gov/edit_collection.html?id=2848), including imaging data from 343 infants in the United States across 812 scans and vocabulary assessments from 185 infants across 382 questionnaires. Following quality control, 222 infants between 10 months and 3 years 6 months with 458 rs-fMRI scans were included in the present analyses. The cohort was rigorously screened: exclusion criteria included preterm birth (< 37 weeks), low birth weight (< 2000 g), major perinatal complications (neonatal hypoxia, NICU stay > 2 days), adoption, first-degree family history of psychiatric disorders (autism, intellectual disability, schizophrenia or bipolar disorder), significant medical or genetic conditions, MRI contraindications, maternal pre-eclampsia, placental abruption, HIV positivity, substance use during pregnancy, and inability of the caregiver to consent in English. Among these, MCDI vocabulary data were available within 0.5, 1, or 2 years of the scan for 111 (214 scans), 138 (293 scans), and 140 (318 scans) participants, respectively. The demographics of subjects with available vocabulary data are described in Table S1.

#### 2. HCP-D

Data were derived from the HCP-Development v2.0 release, a cross-sectional cohort of 652 typically developing children in the United States. Exclusion criteria included premature birth (<37 weeks gestation), serious neurological conditions (e.g., stroke, cerebral palsy), serious endocrine conditions (e.g., precocious puberty, untreated growth hormone deficiency), long-term use of immunosuppressants or steroids, history of serious head injury, hospitalization >2 days for certain physical or psychiatric conditions or substance use, treatment >12 months for psychiatric conditions, claustrophobia, pregnancy, or other contraindications for MRI. We focused on individuals under the age of 18 to capture an age range with documented developmental changes in language acquisition^10,11^. After quality control (see the following section, ‘fMRI Image Quality and Exclusion Criteria,’ for details), 437 participants were included in the analysis.

### fMRI Image Acquisition

#### 1. BCP

All MRI data were acquired on 3T Siemens Prisma scanners with 32-channel head coils at the University of North Carolina at Chapel Hill and the University of Minnesota ^46^. T1-weighted images were obtained using a 3D-MPRAGE sequence (TR/TE/TI = 2400/2.24/1600 ms; flip angle = 8°; 320 × 320 acquisition matrix; 0.8 mm isotropic resolution; 208 sagittal slices). T2-weighted images were acquired via turbo spin-echo (TR/TE = 3200/564 ms; turbo factor 314; echo train length 1166 ms; 0.8 mm isotropic resolution; 208 sagittal slices). Resting-state fMRI utilized a gradient-echo EPI sequence (TR/TE = 800/37 ms; flip angle = 80°; field of view = 208 × 208 mm; 72 axial slices; 2 mm isotropic voxels; 420 volumes), with additional single-band reference and AP/PA reverse-phase scans.

#### 2. HCPD

Imaging was performed on 3T Siemens Prisma systems using protocols harmonized with HCP standards ^47,48^. Structural images included one T1-weighted (MPRAGE; TR/TE/TI = 2500/1.81/1000 ms; flip angle = 8°; 0.8 mm isotropic resolution) and one T2-weighted scan (TR/TE = 3200/564 ms; 0.8 mm isotropic resolution). Resting-state data were collected in two gradient-echo EPI runs (TR/TE = 800/37 ms; 2 mm isotropic voxels; one AP and one PA phase-encoding direction), each lasting 6.5 minutes.

### fMRI Image Processing

#### 1. BCP

Preprocessed data were made available as part of the publication process by Li et al. (2024)^34^. In short, preprocessing was implemented using AFNI (v17.0.08) ^49^ and FSL (v6.0.1) ^50^. Functional volumes were truncated by discarding the first 10 volumes, reoriented, and motion-corrected by alignment to the single-band EPI reference image (SBRef). Distortion correction leveraged AP/PA pairs. Functional data were co-registered with structural images (T1 for those older than 6 months; T2 for younger), then normalized to age-specific templates using ANTs SyN, followed by alignment to a common six-month template via pairwise deformation minimization. Functional data were resampled to 2 mm isotropic resolution, spatially smoothed (4 mm FWHM), detrended, and subjected to nuisance regression including 24 motion parameters, white matter, cerebrospinal fluid, global signal, and regressors for volumes with framewise displacement > 0.5 mm, followed by temporal band-pass filtering (0.01–0.1 Hz).

#### 2. HCPD

This study used the fully preprocessed structural and resting-state fMRI data from the HCP-Development v2.0 release. Details of the preprocessing pipelines are provided in Glasser et al. (2013)^47^. Briefly, structural MRI preprocessing included gradient distortion correction, readout distortion removal, repetition averaging, and nonlinear registration to MNI standard space. Following structural preprocessing, volumetric rs-fMRI data underwent preprocessing, which involved gradient distortion correction, head motion correction, EPI distortion correction, ICA-FIX denoising, registration to both native and MNI standard spaces, and global intensity normalization.

### fMRI Image Quality and Exclusion Criteria

#### 1. BCP

Of the initial 587 resting-state fMRI scans from 292 infants, 8 were removed due to incomplete volumes (< 420), 60 were excluded for lacking structural matches, 42 for excessive motion (mean FD > 0.5 mm and > 40 % high-motion volumes), and 19 for registration or preprocessing failures on visual inspection. The resulting dataset comprised 458 high-quality scans from 222 infants. Among these, MCDI vocabulary data were available within 0.5, 1, or 2 years of the scan for 111 (214 scans), 138 (293 scans), and 140 (318 scans) participants, respectively. The demographics of subjects with available vocabulary data are described in Table S1.

#### 2. HCPD

All individuals included in this dataset had motion below 0.5 mm threshold.

### Cortical and thalamic parcellations

Resting-state BOLD images from all three datasets were parcellated into 400 cortical regions based on the Schaefer 400 atlas ^51^. To identify language-related cortical parcels, the Fedorenko language regions^52^ (https://www.evlab.mit.edu/resources-all/download-parcels), identified with task-based fMRI, were overlaid onto the Schaefer 400 parcellation. For thalamic parcellation in the BCP dataset, the Thalamus Optimized Multi Atlas Segmentation (THOMAS)^53^ was nonlinearly warped to the 6-month template space and subsequently applied to define thalamic subregions.

### Hurst exponent

For this study, we employed the fractional integration method to ensure robust and stable estimation. fractional integration models treat the time series as fractionally integrated processes, where the Hurst exponent H is defined as H = *d* + 0.5, with *d* representing the fractional integration order ^42,54,55^. In these models, the autocorrelation function (ACF) decays according to a power law, approximately as ACF(*k*) ∼ *k*^2d−1^, where *k* is the lag. This means that for *d* > 0, the correlations between distant observations decline slowly, reflecting long-range dependence or persistence in the signal. This approach fits parametric models to capture long-range dependencies more rigorously. In this study, the Hurst exponent was computed using the *nonfractal* MATLAB toolbox (https://github.com/wonsang/nonfractal)^42,54^. The specific function utilized is bfn_mfin_ml.m function with the ‘filter’ argument set to ‘haar’ and the ‘ub’ and ‘lb’ arguments set to [1.5,10] and [−0.5,0], respectively. This toolbox uses a discrete wavelet transform and a model of the time series as fractionally integrated processes and is estimated using maximum likelihood estimation..

### Global cortical hierarchies

Because the secondary functional cortical hierarchy in infants has been shown to be an anterior-posterior gradient^33^, we followed the methods of Lariviere and colleagues^33^ to calculate the A-P gradient using the BCP dataset. First, we calculated pairwise correlations between all 400 cortical parcels to construct a functional connectome. The resulting correlation matrix was z-transformed and averaged to produce a group-level connectome. We retained only the top 10% of connections and transformed the thresholded connectivity matrix into a normalized angle matrix, which scales the angle between vertex pairs according to their similarity. We then applied diffusion map embedding^56,57^. The resulting gradient closely matched the map published by Lariviere and colleagues^33^.

The sensorimotor-association (S-A) axis was derived by Sydnor and colleagues^48^.^58^ This map encompasses various cortical hierarchies, including functional connectivity gradients, evolutionary cortical expansion patterns, anatomical ratios, allometric scaling, brain metabolism measures, perfusion indices, gene expression patterns, primary modes of brain function, cytoarchitectural similarity gradients, and cortical thickness.

To assess the relationship between each cortical gradient and age-related effects on the Hurst exponent, we used Spearman’s rank correlation and evaluated statistical significance using spin-based spatial permutation tests ^59,60^, which preserve the spatial covariance structure typical in neuroimaging data. A null distribution was generated through 10,000 spherical rotations, against which the observed correlation was compared.

### Demographics and Socioeconomic Status

For the BCP dataset, parental education and household income were included as measures of socioeconomic status (SES). When education levels for both parents were available, their average was used; if data were available for only one parent, that value was used. In the BCP dataset, participants were classified as bilingual if the primary language spoken at home accounted for less than 90% of household language use. For the HCP-D dataset, only household income was available.

### Language outcomes

In the BCP dataset, vocabulary was assessed using the MacArthur-Bates Communicative Development Inventories (MCDI)^61^, focusing on two specific measures: total word comprehension and total word production. These measures involve questions such as “Does your child understand the word ‘ball’?” or “Does your child say the word ‘mama’?”, where caregivers report whether the child understands or produces specific words from a standardized vocabulary list. BCP participants completed the MCDI at one to four timepoints. Because the timing of the MCDI assessments and brain scans differed, we calculated the time difference between each scan and the corresponding questionnaire (referred to as the age gap), and limited our analysis to age gaps of 0.5, 1, or 2 years to ensure temporal proximity.

### Statistical models

To flexibly model both linear and nonlinear relationships between the Hurst exponent and age, we used Generalized Additive Models (GAMs) implemented via the *mgcv* package in R. In these models, the region-averaged Hurst exponent served as the dependent variable, age was included as a smooth term using a penalized regression spline with a maximum basis dimension of *k* = 3, and sex and mean framewise displacement were included as linear covariates. For the BCP dataset, we also controlled for site effects. Additionally, a random intercept for participant ID was included to account for within-subject variability. This factor-smooth interaction models subject-specific deviations from the population-level smooth, thereby accommodating the longitudinal structure of the data and appropriately capturing correlations among repeated measures. Models were fitted separately for each parcellated cortical region. The smooth term for age generated a spline representing a region’s developmental trajectory.

For each regional GAM, the significance of the smooth term for age was assessed using an approximate F-test, as implemented in *mgcv*. This test evaluates whether the coefficients of the spline basis functions for the smooth are all equal to zero, while accounting for the estimated effective degrees of freedom of the smooth. A significant F-statistic indicates that the smooth explains variance in the Hurst exponent beyond the intercept-only model. For each regional GAM, we identified the specific age range(s) where the Hurst exponent significantly changed using the *gratia* package in R. Age windows of significant change were determined by examining the first derivative of the age smooth function (Δ Hurst exponent/Δ age) and assessing when the simultaneous 95% confidence interval of this derivative did not include 0 (two-sided).

To investigate interactions between age and language skill, we used GLMs. The region-averaged Hurst exponent was used as the dependent variable. Age-by-language interaction, sex, mean framewise displacement, household income, parental education, and site were included as linear covariates. For each regional GLM, we evaluated the significance of interaction effects using t-statistics for the regression coefficients and their associated p-values. False Discovery Rate (FDR) correction was applied across the six language ROIs to control for multiple comparisons.

## Results

### Hurst in language-related cortex increases across early childhood

To examine the development of neural inhibition in language-related brain regions in infancy, we first analyzed the Hurst exponent of resting-state fMRI data from the Baby Connectome Project (BCP). We observed significant positive associations between age and the Hurst exponent across both temporal and frontal language regions across the age range (Figure 1A, Temp1: *R*^2^ = 0.005, *P*_FDR_ < 0.001; Temp2: *R*^2^ = 0.013, *P*_FDR_ < 0.001; Temp3: *R*^2^ = 0.022, *P*_FDR_ < 0.001, Figure 1B, Front1: *R*^2^ = 0.003, *P*_FDR_ = 0.002; Front2: *R*^2^ = 0.003, *P*_FDR_ < 0.001; Front3: *R*^2^ = 0.009, *P*_FDR_ < 0.001). The age effect was especially pronounced in the posterior temporal language regions (Figure 1C). The most posterior frontal region exhibited the strongest age effect, showing a marked increase from age 3 to age 3.5 (Figure 1C).

**Figure 1.**
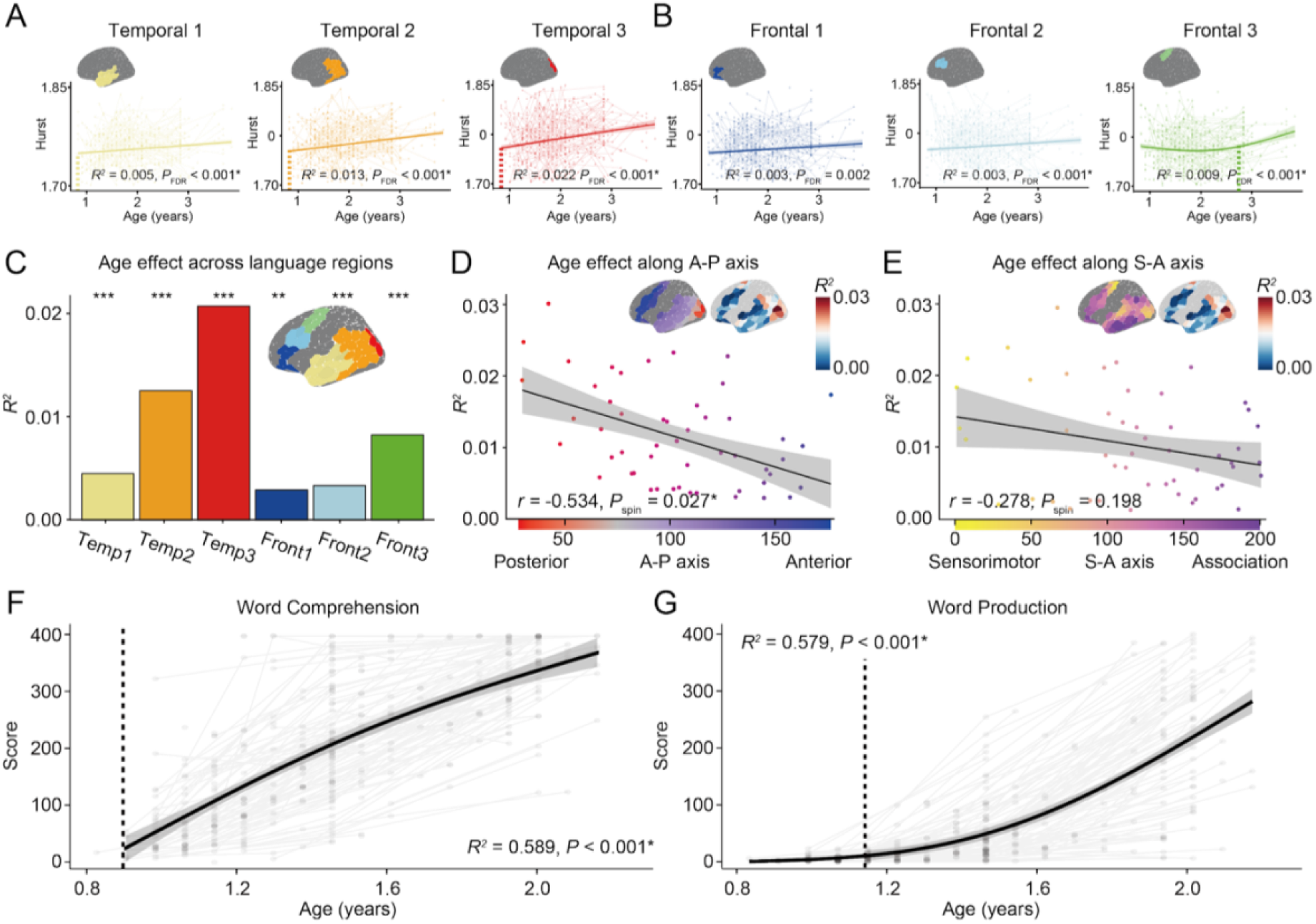
Age effects across cortical language regions in infancy. **(A, B)** Developmental trajectory of the Hurst exponent in temporal (A, Temp1: *R*^2^ = 0.005, *P*_FDR_ < 0.001; Temp2: *R*^2^ = 0.013, *P*_FDR_ < 0.001; Temp3: *R*^2^ = 0.022, *P*_FDR_ < 0.001) and frontal (B, Front1: *R*^2^ = 0.003, *P*_FDR_ = 0.002; Front2: *R*^2^ = 0.003, *P*_FDR_ < 0.001; Front3: *R*^2^ = 0.009, *P*_FDR_ < 0.001) regions. Motion, sex, and site effects were regressed out, and the mean Hurst value was added back to preserve the average for each ROI. The shaded areas represent the 95% confidence intervals. The dashed line marks the age at which the first derivative of the age smooth function (Δ Hurst exponent / Δ age) becomes significant. **(C)** Age effects across temporal and frontal language regions. \**P* < 0.05, \*\**P* < 0.01, \*\*\**P* < 0.001. **(D, E)** Age effects of language regions along anterior-posterior (A-P) axis (D, *r* = −0.534, *P*_spin_ = 0.027) and sensorimotor-association (S-A) axis (E, *r* = −0.278, *P*_spin_ = 0.198). The shaded areas represent the 95% confidence intervals. The supplementary image above the graph shows the R² for each parcel (low to high, blue to red). **(F, G)** Developmental trajectory of word comprehension (F, *R*^2^ = 0.589, *P* < 0.001) and word production (G, *R*^2^ = 0.579, *P* < 001). The shaded areas represent the 95% confidence intervals. The dashed line marks the age at which the first derivative of the age smooth function (Δ Word score / Δ age) becomes significant.

### Hurst development is graded along the anterior-posterior (A-P) axis

Given that global cortical maturation follows hierarchical patterns^58^, we tested whether age effects across language regions align with these macroscale axes. Regional differences in age effects across language regions were well explained by the A-P axis (Figure 1D, *r* = −0.534, *P*_spin_ = 0.027), whereas the S-A axis did not account for the variance (Figure 1E, *r* = −0.278, *P*_spin_ = 0.198).

### Trajectories of growth in language comprehension and production parallel Hurst development

We next turned to behavioral measures to explore how the neural development of Hurst related to the development of language-related skills. Word comprehension scores increased rapidly before slowing down around 18 months of age (Figure 1F, *R*^2^ = 0.589, *P* < 0.001), whereas word production scores began to rise around 14 months of age (Figure 1G, *R*^2^ = 0.579, *P* < 0.001), consistent with previously established norms^62,63^.

### Protracted development of the Hurst exponent is linked to advanced vocabulary

Given the alignment between the developmental trajectories of language skills and the Hurst exponent, we investigated whether an individual-level association exists between the two. The scan and questionnaire timings for each participant are shown in Figure S1. Our primary focus was on whether language abilities moderated age-related changes in the Hurst exponent. Because this analysis could take many different specifications (time between scan and behavioral measure, language measure, control variables), we used a specification curve analysis to test all possible effects. The specification curve from our multiverse analysis revealed a consistent negative interaction, suggesting that higher language skills are associated with a slower developmental increase in the Hurst exponent (Figure 2A). This interaction was significant primarily in temporal, but not frontal, language regions. Notably, effect sizes were larger when the age gap was smaller (i.e., 0.5 or 1 year), despite the reduced sample size (age gap 0.5 year; *N* = 111, *scans* = 214, 1 year; *N* = 138, *scans* = 293, 2 years; *N* = 140, *scans* = 318). This may be due to better alignment between brain and behavior at closer timepoints. Furthermore, the interaction effects tended to be stronger for language comprehension than for production, likely due to the stronger coupling between the temporal regions and language comprehension. Controlling for SES had minimal impact on the overall pattern of results.

**Figure 2.**
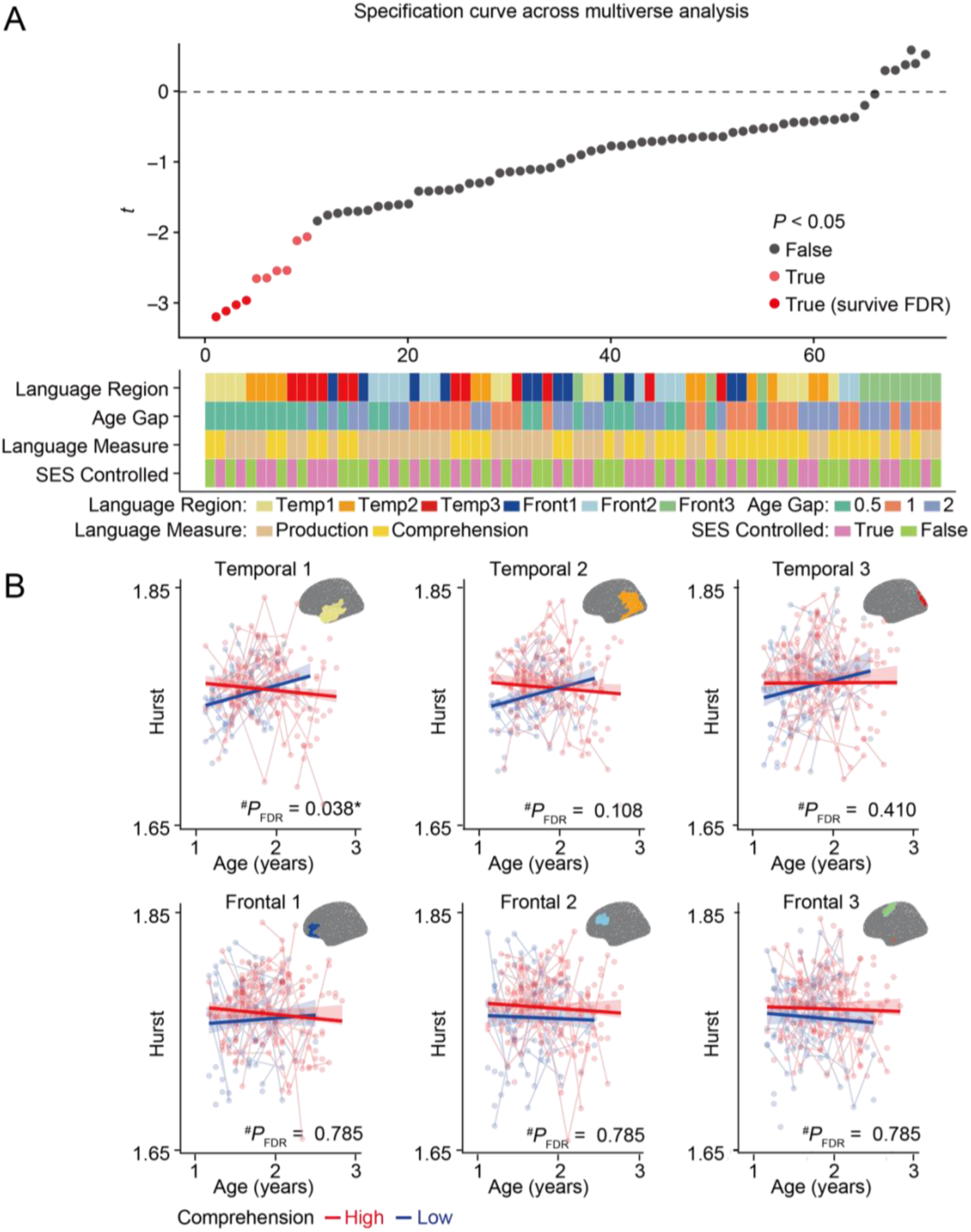
Interactions between the Hurst exponent and language development. **(A)** Each point on the specification curve represents a specific analytical model, varying by language region, age gap threshold (0.5, 1, or 2 years), language measure (comprehension vs. production), and inclusion of socioeconomic status (SES) as a covariate. The y-axis displays the estimated interaction t-statistics, while the x-axis ranks the models by effect size. Negative values indicate that higher language skills are associated with a slower developmental increase in the Hurst exponent. Dark red dots indicate models where the interaction effect is statistically significant (FDR-corrected across six language regions), while light red dots do not survive FDR correction. **(B)** Visualization of developmental trajectories of the Hurst exponent in children with high (red) versus low (blue) word comprehension skills (Temp1 *P*_FDR_ = 0.038, Temp2 *P*_FDR_ = 0.108, Temp3 *P*_FDR_ = 0.410, Front1 *P*_FDR_ = 0.785, Front2 *P*_FDR_ = 0.785, Front3 *P*_FDR_ = 0.785). This model uses an age gap of 0.5 years, the comprehension measure, and includes SES covariates. Motion, sex, and site effects were regressed out, and the mean Hurst value was added back to preserve the average for each ROI. The shaded areas represent the 95% confidence intervals.

For visualization purposes only, we selected the specification that showed the strongest age × comprehension interaction (age gap = 0.5 years, comprehension measure, SES controlled) and plotted the developmental trajectories of the Hurst exponent for children with high versus low language comprehension (above or below the mean; Fig. 2B). This choice was made post hoc for interpretability and does not form the basis of statistical inference. After applying FDR correction across the six language regions, the interaction effect remained statistically significant in the most anterior temporal region (Temp1: *P*_FDR_ = 0.038, Temp2: *P*_FDR_ = 0.108, Temp3: *P*_FDR_ = 0.410, Front1: *P*_FDR_ = 0.785, Front2: *P*_FDR_ = 0.785, Front3: *P*_FDR_ = 0.785). This interaction effect remained significant even after controlling for bilingualism, despite the limited sample size due to bilingualism information being available from only one of the two data collection sites (*N* = 92, *scans* = 189, Temp1: *P*_FDR_ = 0.046).

### Thalamic maturation precedes cortical maturation

Given that we found that cortical Hurst was still increasing after the first year of life, we investigated whether earlier language sensitive periods might align instead with the development of thalamic nuclei. Previous studies suggest that the sensitive period for phoneme perception occurs by 12 months.^2,3^ We examined the development of Hurst in the medial geniculate nucleus (MGN) of the thalamus, which supports language comprehension^26–28^, and the ventrolateral anterior and posterior (VLa/VLp) nuclei, which are involved in language production^29,30^. To test the hypothesis that thalamic neural inhibition matures earlier than cortical inhibition, we examined the developmental trajectories of the Hurst exponent in MGN and VLa and VLp. We applied FDR correction across three nuclei. The age effect survived FDR correction only in VLp. MGN showed a linear decreasing pattern (Figure 3A, *R*^2^ = 0.010, *P*_FDR_ = 0.112), while VLa exhibited a slight non-linear pattern, with a decline beginning around age 2 (Figure 3B, *R*^2^ = 0.012, *P*_FDR_ = 0.111). In VLp, we observed an increase until age 2 followed by a decline (Figure 3C, *R*^2^ = 0.050, *P* < 0.001, *P*_FDR_ < 0.001). These results are consistent with the theory that thalamic nuclei reach their developmental plateau in neural inhibition earlier than cortical language regions, timing which may parallel the sensitive period for phoneme perception.

**Figure 3.**
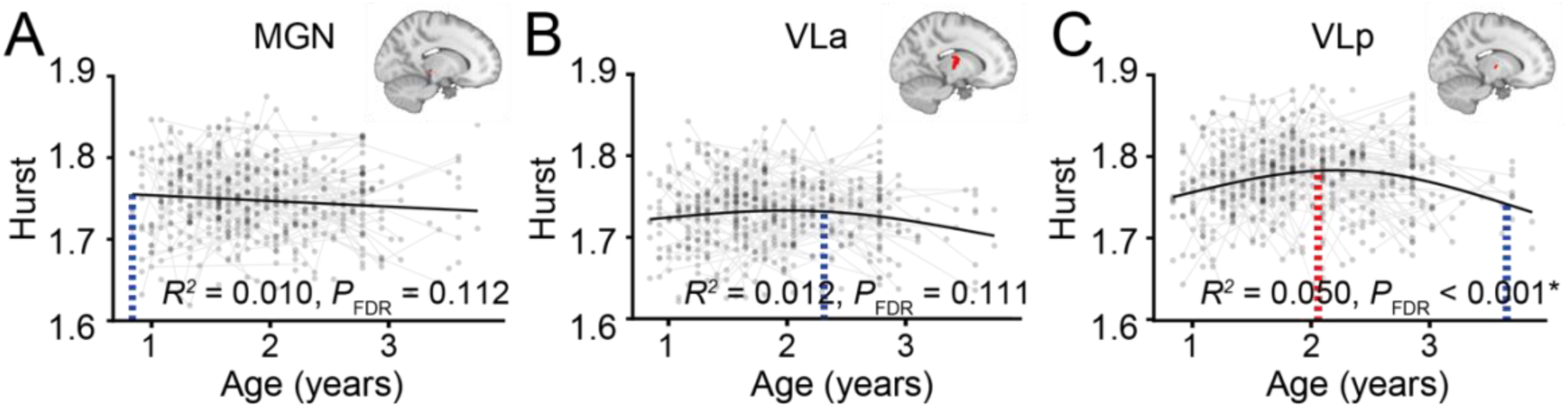
Age effects across language-related thalamic nuclei during infancy. **(A)** Developmental trajectory of the Hurst exponent in MGN (*R*^2^ = 0.010, *P*FDR = 0.112). **(B)** Developmental trajectory of the Hurst exponent in VLa (*R*^2^ = 0.012, *P*FDR = 0.111, *Decrease Age* = 2.35 y). **(C)** Developmental trajectory of the Hurst exponent in VLp (*R*^2^ = 0.050, *P*FDR < 0.001, *Increase Age* = 2.05 y, *Decrease Age* = 3.62 y). The dashed line indicates the age at which the first derivative of the age smooth function (Δ Hurst exponent / Δ age) becomes significant (blue: negative, red: positive). Motion, sex, and site effects were regressed out.

### The Hurst exponent reaches a plateau in middle childhood along the sensorimotor - association axis

Prior work suggests a sensitive period for syntax learning around ages 8–10^8,9,64^. To test whether the maturation of neural inhibition in language regions corresponds to the decline in children’s ability to learn new syntax during childhood, we analyzed the Hurst exponent in children aged 6 to 18 years (*N* = 437) using the Human Connectome Project–Development dataset. We observed a continuous increase in the Hurst exponent during childhood, with a plateau emerging in late childhood around 8 to 10 years of age, corresponding to the proposed sensitive period for syntax. The frontal regions showed a slightly later plateau compared to the temporal regions (Temp1; *R*^2^ = 0.007, *P*_FDR_ < 0.001, *Age* = 6.33 y, Temp2; *R*^2^ = 0.005, *P*_FDR_ < 0.001, *Age* = 9.05 y, Temp3; *R*^2^ = 0.004, *P*_FDR_ = 0.004, Front1; *R*^2^ = 0.013, *P*_FDR_ < 0.001, *Age* = 10.41 y, Front2; *R*^2^ = 0.014, *P*_FDR_ < 0.001*, Age* = 10.29 y, Front3; *R*^2^ = 0.020, *P*_FDR_ < 0.001, *Age* = 9.91 y). In contrast to infancy, age-related effects were stronger in frontal language regions than in temporal regions during this period (Figures 4B). Unlike in infancy, the S-A axis, rather than the A-P axis, accounts for age-related changes during this period, with association regions showing larger age effects. This suggests a “catch-up” of regions that exhibited weak age effects in infancy (AP; Figure 4C, *r* = 0.316, *P*_spin_ = 0.171, SA; Figure 4D, *r* = 0.520, *P*_spin_ < 0.001).

**Figure 4.**
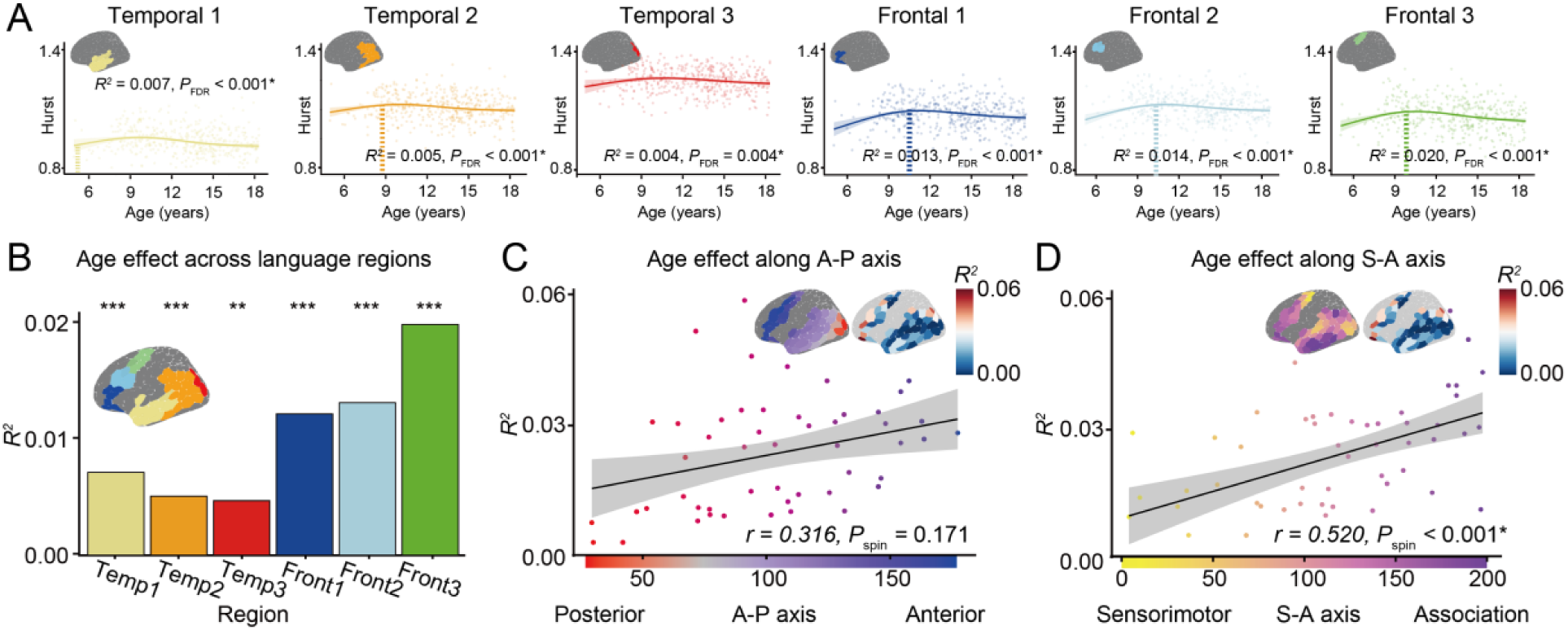
Age effects across cortical language regions in childhood. **(A)** Developmental trajectory of the Hurst exponent across language regions. Motion, sex, and site effects were regressed out, and the mean Hurst value was added back to preserve the average for each ROI. The shaded areas represent the 95% confidence intervals (Temp1; *R*^2^ = 0.007, *P*_FDR_ < 0.001, *Age* = 6.33 y, Temp2; *R*^2^ = 0.005, *P*_FDR_ < 0.001, *Age* = 9.05 y, Temp3; *R*^2^ = 0.004, *P*_FDR_ = 0.004, Front1; *R*^2^ = 0.013, *P*_FDR_ < 0.001, *Age* = 10.41 y, Front2; *R*^2^ = 0.014, *P*_FDR_ < 0.001*, Age* = 10.29 y, Front3; *R*^2^ = 0.020, *P*_FDR_ < 0.001, *Age* = 9.91 y). The dashed line marks the age at which the first derivative of the age smooth function (Δ Hurst exponent / Δ age) becomes insignificant. **(B)** Age effects across temporal and frontal language regions. \**P* < 0.05, \*\**P* < 0.01, \*\*\**P* < 0.001. **(C, D)** Age effects of language regions along anterior-posterior (A-P) axis (C, *r* = 0.316, *P*_spin_ = 0.171) and (D, *r* = 0.520, *P*_spin_ < 0.001) sensorimotor-association (S-A) axis. The shaded areas represent the 95% confidence intervals. The supplementary image above the graph shows the *R*² for each parcel (low to high, blue to red).

## Discussion

Here, we explored the neural basis for sensitive periods in language development. We used the Hurst exponent to characterize neural inhibition, a key modulator of neuroplasticity, in language-related brain regions from infancy to late childhood. We found that inhibition increased in temporal and frontal language-related cortex across early childhood. This increase in inhibition was not uniform across language cortex, but rather graded along cortical hierarchies. In contrast to the pattern in cortex, Hurst in thalamic nuclei associated with auditory processing and motor control plateaued within the first three years of life. This more closely aligns with the timing of the earliest language-related sensitive periods, including declines in non-native phoneme sensitivity. Thus, our data raise questions about whether this initial sensitive period is instantiated in thalamus, rather than cortex. In contrast, we found that cortical Hurst ultimately plateaued between ages 8-10, paralleling evidence for a potential sensitive period in syntax learning around this age. Finally, we found some evidence that young children with higher early vocabularies had later-occurring increases in inhibition in language-related cortex. This suggests there may be prolonged plasticity for those with the highest language-related skills.

We found that Hurst increased across temporal and frontal language-related cortex in the first few years of life. This pattern of data suggests that language-related sensitive periods instantiated in the cortex remain open past the first few years of life, contrary to our initial predictions. Thus, the earliest language-related sensitive periods in language do not appear to be instantiated in cortex. The trajectories of Hurst in temporal language regions followed similar developmental patterns as children’s developing comprehension skills, while trajectories of Hurst in frontal language regions more closely paralleled children’s developing production skills. However, more research is needed to point to a mechanistic link between Hurst development and children’s developing language-related skills.

We also found that the maturation of Hurst in temporal and frontal language-related cortex was not uniform, but rather graded across well-known hierarchies of brain development. At the youngest ages, Hurst increased along the anterior-posterior axis, such that inhibition began to increase later for more anterior regions. Thus, in general, the more posterior temporal areas matured faster than the more anterior frontal areas. This hierarchical gradient may be a precursor to the sensorimotor-association (S-A) axis, which shows graded development across later childhood.^31,33,34^ Indeed, among older children in our sample, the S-A axis better explained gradations in Hurst. This natural hierarchy may underly the graded nature of the development of language-related skills^1,31^. Language comprehension emerges earlier than production; in our sample, the time course of these developing skills visually mapped onto the development of inhibition for temporal and frontal language-related cortex, respectively.

Our results suggest that language-related thalamic nuclei showed earlier maturation than cortex. Indeed, the thalamic nuclei we examined showed patterns of Hurst that paralleled the timing of early language-related sensitive periods, with inhibition plateauing in the first year of life. The medial geniculate nucleus (MGN) has connections to the auditory cortex and has been shown to play a role in processing phoneme distinctions,^24–28^ and appeared to have already reached a plateau in Hurst by age 10 months. This adds to growing evidence for the possibility that the sensitive period for phonemic discrimination—the ability to hear non-native speech sounds—may result from maturation of thalamus rather than cortex. Indeed, longstanding work points to the role of the MGN in processing speech sounds and gating information to the cortex, leading to arguments that the thalamus gates the discrimination of non-native speech phonemes^28,65^.

We found that on average, Hurst in language-related cortex plateaued between ages 8-10 years old. The timing of this plateau aligns with longstanding evidence that a second language becomes harder to learn after these ages.^8,9,66^ More recently, scholars have argued that the sensitive period for learning grammar in a new language extends until 18 years, and that prior findings had to do with years of exposure.^10^ While we don’t have behavioral data to bear on this question, the timing of the plateau in Hurst in our sample aligns best with the former suggestion. These declines in plasticity may provide some neural evidence as to why there are declines in second language learning around age 9-10 years.

In terms of individual differences, we found that higher vocabulary was linked to protracted development of temporal language regions. Between 10 months and 3.5 years, we found a significant interaction, such that children with lower vocabulary comprehension and production had steeper, more positive slopes of Hurst, suggesting that plasticity was declining more rapidly. Given that children with earlier phonetic commitment to their native language show more rapid vocabulary growth, one possibility we considered was that higher vocabulary would be associated with earlier sensitive periods. Indeed, some theories have suggested that plasticity may close when a child has received sufficient input to commit to a language, independent of other factors.^67^ However, our findings argue against this possibility. Because children with the highest vocabularies in infancy are those who receive the most linguistic input^68^, our findings are more in line with theories that richer, more dynamic input may lead to protracted sensitive periods.^1^ This more dynamic input extending sensitive periods may also help to explain why children learning multiple languages may show more protracted sensitive periods.^67,69^ It may also help explain recent findings that young children with high phonological processing skills show more long-lasting increases in the volume and surface area of several language-related brain regions.^70^ Of course, it will be important in future work to measure these neural developmental trajectories in tandem with behavioral measures of phonemic discrimination.

Our work raises questions about whether protracted periods of sensitivity are due to language input or other variables such as stress or other features of the child’s environment. In our study, the results held when controlling for children’s family socioeconomic status. Children’s language input is related to their language learning above and beyond SES^71^, whereas effects of stressful life events appear to be mediated through language input^72^; thus, we believe our findings point to a more direct role for a child’s language input. Still, it is possible that children whose vocabularies were lower also experienced more stressful life events, which sped up their neural development.^43^ The children in our samples also came from relatively high-socioeconomic backgrounds. In the BCP dataset, nearly three-quarters of caregivers had a college degree or higher, while in the United States in 2022, about 37.7% of adults age 25 and over had a college degree or higher^61^.^73^ In the HCPD dataset, in which parental education was not available, 60% of families had incomes above $100K/year, while in the United States in 2022, approximately 37.8% of families had incomes above $100k.^74^ Thus, future work should examine plasticity in language-related brain regions for children from a more diverse range of backgrounds and measure children’s stress and language input directly.

Several additional potential limitations of this study should be acknowledged. First, our measures of children’s language development were limited. We only had parent-reported measures of infants’ word knowledge as opposed to direct assessments, and no tests that measured children’s phoneme distinction. Moreover, the BCP collects brain imaging and vocabulary measures at different timepoints, which limits the temporal precision with which neural and behavioral data can be linked. Second, in the BCP dataset, infants under 3 years old were sleeping during the scan, whereas scans from children older than 3 years included a mix of sleeping and awake states. Prior work on validation of the Hurst exponent was done with awake resting-state data so we do not know how sleep data might be different. Third, data were preprocessed for the two studies by independent groups who made some different choices. Differences in nuisance regression strategies (e.g., global signal regression in the BCP dataset but not in the HCPD dataset) and motion censoring procedures (scrubbing with FD > 0.5mm for the BCP dataset but ICA-FIX denoising for the HCPD dataset) may contribute to differences in absolute values of the Hurst exponent across datasets, but are unlikely to impact developmental trajectories or individual differences within dataset.

In sum, this work elucidated trajectories of inhibitory development for language-related brain regions across cortex and thalamus. Our findings point to graded periods of plasticity across language-related brain regions, perhaps driven by global cortical hierarchies. It is possible that these cortical hierarchies underlie the graded nature of language-related sensitive periods that have been observed behaviorally, such that, for example, language perception skills develop earlier than language production skills. Our data also add to growing evidence suggesting the earliest-observed language-related sensitive periods may be instantiated in thalamus, rather than cortex, though more work is needed to measure this directly. Ultimately, understanding these sensitive periods may be useful for time-sensitive language-related interventions, as well as for our basic understanding of the basis of the human language system.

## Data and code availability

Data used in this study were obtained from the Baby Connectome Project (BCP) and the Human Connectome Project–Development (HCP-D). Both datasets are publicly available through the NIMH Data Archive (NDA; https://nda.nih.gov/). Access to each dataset requires an approved Data Use Certification. All original code is available at [removed for peer review].

## Acknowledgements

We are grateful to Jed Ellison, Lana Hantzsch, Qingling Li, and Yong He for their help with data procurement. This work was funded by National Institute of Mental Health Project Number 2R01MH113550 to TS and APM and National Science Foundation Award #2045095 to APM.

## Conflicts of Interest

The authors have no conflicts of interests to disclose.

## Notes

https://github.com/monami-nishio/hurst_language_plasticity

### Competing Interest Statement

The authors have declared no competing interest.

### Summary of Updates

The current version of the manuscript updates one formatting error.

