## Supplementary Materials for "Cascading periods of language-related brain plasticity across early childhood"

Monica E. Ellwood-Lowe*[1], Monami Nishio*[2], Alexander J. Dufford[3],

Michael Arcaro[2], Theodore D. Satterthwaite[4,5,6], Allyson P. Mackey[2]

### Supplement

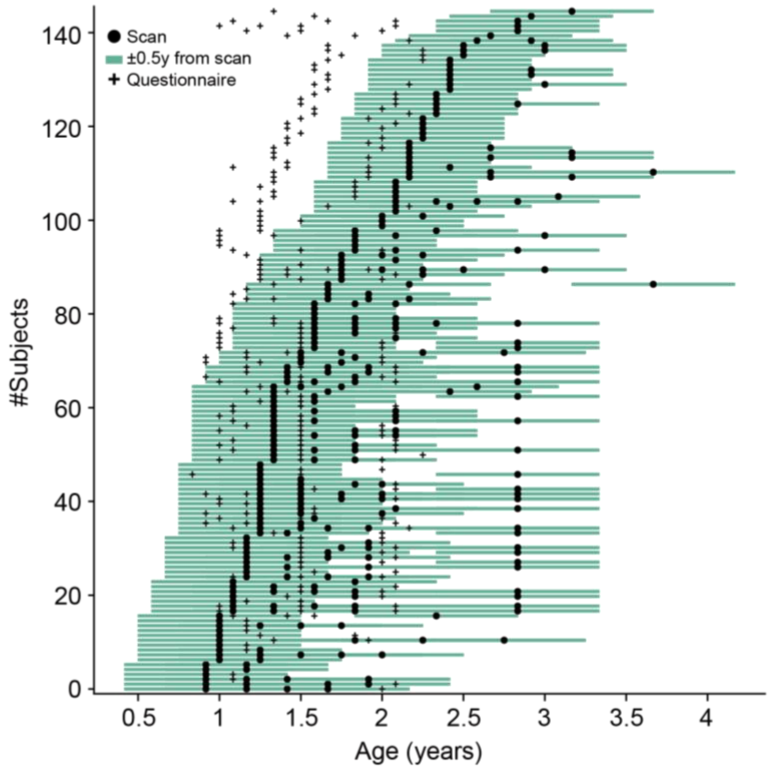


**Supplemental Figure 1. Timing of scans and questionnaires for each participant in BCP.** The timing of the scan (circle) and the language questionnaire (cross) is shown for each participant. Participants are ordered by the timing of their scan. The green shaded area represents the ± 0.5 year window around the scan date.

### Table S1. Demographics of BCP subjects with vocabulary data.

| | **±0.5y (*N* = 111) mean or #** | **±1y (*N* = 138) mean or #** | **±2y (*N* = 140) mean or #** |
|---|---|---|---|
| **Age** (years) | 1.599 ± 0.3997 | 1.807 ± 0.5325 | 1.889 ± 0.5903 |
| **Sex** (male) | 53 (48%) | 66 (48%) | 66 (47%) |
| **Ethnicity** | | | |
| Hispanic or Latino | 7 (6%) | 11 (8%) | 11 (8%) |
| No Response | 0 (0%) | 0 (0%) | 0 (0%) |
| **Race** | | | |

| | | | |
|---|---|---|---|
| White | 89 (80%) | 111 (81%) | 113 (81%) |
| Black | 1 (1%) | 2 (1%) | 2 (1%) |
| Asian | 0 (0%) | 1 (0%) | 1 (0%) |
| Others | 21 (19%) | 24 (18%) | 24 (18%) |
| No Response | 0 (0%) | 0 (0%) | 0 (0%) |
| **Income** | | | |
| 25-35k | 4 (4%) | 6 (4%) | 6 (4%) |
| 35-50k | 3 (3%) | 3 (2%) | 3 (2%) |
| 50-75k | 20 (18%) | 26 (19%) | 27 (19%) |
| 75-100k | 21 (19%) | 24 (17%) | 24 (17%) |
| 100-150k | 36 (32%) | 45 (33%) | 45 (32%) |
| 150-200k | 17 (15%) | 21 (15%) | 22 (16%) |
| 200k- | 9 (8%) | 12 (9%) | 12 (9%) |
| No Response | 1 (0%) | 1 (0%) | 1 (0%) |
| **Parent Education** | | | |
| Less than college | 7 (6%) | 9 (7%) | 9 (6%) |
| College degree | 84 (77%) | 100 (73%) | 102 (74%) |
| Graduate degree | 19 (17%) | 28 (20%) | 28 (20%) |
| No Response | 1 (0%) | 1 (0%) | 1 (0%) |
| **Multilingual** | | | |
| Multilingual | 17 (15%) | 22 (16%) | 22 (16%) |
| Monolingual | 77 (70%) | 92 (67%) | 93 (66%) |
| No Response | 17 (15%) | 24 (17%) | 25 (18%) |
